# Influence of visual information on sniffing behavior in a routinely trichromatic primate

**DOI:** 10.1101/2023.05.24.542038

**Authors:** Brigitte M. Weiß, Anja Widdig

## Abstract

Most catarrhine primates are considered to be strongly visually oriented, obtaining information about conspecifics and their environment from a diversity of visual cues. Other sensory modalities may provide information that is redundant and/or complimentary to visual cues. When cues from multiple sensory modalities are available, these may reinforce or suppress each other, as shown in several taxa ranging from insects to humans. Here, we tested how the presence and ambiguity of visual information affects the use of olfactory cues when exploring food and non-food items in semi-free ranging Barbary macaques at Affenberg Salem, Germany. We presented monkeys with pipes containing food (peanuts, popcorn), non-food (stones, feces) or no items in transparent or opaque containers, and assessed whether animals looked, sniffed and/or grabbed into the pipes depending on visibility of the contents (experiment 1). Visual information had no robust effect on sniffing probability, but it did affect sniffing behavior with respect to the timing of olfactory inspections (i.e. whether sniffing occurred first). Both visual and olfactory information affected, whether or not monkeys attempted to retrieve the items from the pipes. Furthermore, we manipulated the visual appearance of familiar food items (popcorn) with food colorant (experiment 2), which resulted in substantially increased olfactory inspections compared to unmanipulated popcorn. Taken together, reliance on the olfactory sense was modulated by the available visual information, emphasizing the interplay between different sensory modalities for obtaining information about the environment.

## INTRODUCTION

Animals gain information about their physical and social environment using different sensory channels. Information can be conveyed via vision, audition, olfaction or other sensory modalities either by a single or multiple sensory channels in combination. In fact, sensory information is frequently multi-modal (i.e. encompassing more than one modality) in nature and integrating information from multiple modalities is widespread throughout the animal kingdom (e.g. Rushmore et al. 2012; Higham and Hebets 2013). Well-known examples include aposematically colored insects, which typically also emit an odor or sound when approached by a predator, or courtship signals encompassing combinations of visual, acoustic, tactile and/or chemical elements in a range of taxa (e.g. insects, fish, birds, see review in Hebets and Papaj 2005). The different modalities may thereby provide redundant or complimentary information simultaneously or sequentially, which may reinforce or in other ways interact with each other, and/or act at different spatial or temporal scales (Hebets and Papaj 2005). As a result, multimodal information may be more accurate, salient or diverse, more robust to noise, more memorable or available across a larger range than information conveyed via a single modality (Hebets and Papaj 2005; Higham and Hebets 2013; Leonard and Masek 2014).

Many multimodal studies focus on relatively easily recordable modalities like vision and audition, but an increasing number of studies show multimodal interactions involving olfaction as well (reviewed in Hebets and Papaj 2005). The interplay of olfaction with other senses has been studied relatively well in insects and insect-plant interactions (reviewed for bees in Leonard and Masek 2014), but rather little in vertebrates (see Melin et al. 2019). Nonetheless, there is evidence that the availability and salience of information in one modality may affect to what extent vertebrates attend to other sensory modalities, including olfaction, in various contexts (e.g. communication in lizard *Liolaemus pacha,* Vicente and Halloy 2017; foraging bats *Artibeus watsoni* and *Vampyressa pusilla*, Korine & Kalko 2005; odor detection in humans, Gottfried and Dolan 2003).

Among primates, vision and olfaction have been suggested to be the main senses in foraging ecology (e.g. Dominy 2004; Rushmore et al. 2012; Veilleux et al. 2022). Most primates are strongly visually oriented animals (Kawamura 2016). They have three-dimensional vision and are the only eutherian mammals to have developed trichromatic vision, albeit not in all taxa (Jacobs 2009; Kawamura 2016). More specifically, catarrhine primates (Old World monkeys and apes) and howler monkeys have evolved routine trichromacy, while many platyrrhine species (New World monkeys) show a polymorphism that results in heterozygous females being trichromatic and homozygous females as well as all males being dichromatic (Kawamura 2016). Primates in all taxa also routinely rely on olfaction in foraging, social and sexual contexts (e.g. Drea 2015; Vaglio et al. 2016; Jänig et al. 2018; Kücklich et al. 2019). Importantly, the reliance on olfaction has been suggested to be related to visual capabilities, as dichromatic white-faced capuchins (*Cebus imitator*) sniffed fruits more often than trichromatic conspecifics (Melin et al. 2019). However, no clear relationship between sniffing and chromacy was found in black-handed spider monkeys (*Ateles geoffroyi*, Hiramatsu et al. 2009). Similarly, the sense of smell was reported to have higher subjective value in blind than sighted humans and olfactory bulb size was found to be larger in early blind people (reviewed in Sorokowska et al. 2019), although there is no robust evidence for a behavioral difference in olfactory abilities between blind and sighted people when corrected for publication bias (Sorokowska et al. 2019).

Reliance on olfaction in primates has further been suggested to be related to feeding ecology and the sensory cues provided by food items (Nevo and Heymann 2015). For instance, black-handed spider monkeys (Hiramatsu et al. 2009), but not white-faced capuchins (Melin et al. 2019), were found to sniff more at visually less conspicuous fruits. An interplay between different senses in foraging success was shown in nocturnal grey mouse lemurs (*Microcebus murinus*), which were able to detect their insect prey using either visual, acoustic or olfactory cues in isolation but performed best when cues from all three modalities were available together (Piep et al. 2008). Similarly, folivore Coquerel’s sifakas (*Propithecus coquereli*) required visual and olfactory cues together to reliably identify more nutritious food, while generalist ring-tailed lemurs (*Lemur catta*) could use either modality alone, and frugivore ruffed lemurs (*Varecia variegata*) could use olfactory, but not visual, cues alone to select the more nutritious food items (Rushmore et al. 2012). Hence, current evidence points towards olfaction in primate feeding ecology being relevant primarily for food selection and when visual information about food quality is lacking or ambiguous, although the overall evidence is mixed and data are still too scarce for robust conclusions (Hiramatsu et al. 2009; Nevo and Heymann 2015). Furthermore, with the exception of a few studies in humans, data on the interplay between vision and olfaction are lacking for species that are routinely trichromatic.

Barbary macaques (*Macaca sylvanus*) are feeding generalists (Fooden 2007) with trichromatic vision (Niimura et al. 2018). They use olfaction in a range of contexts but direct the majority (∼80%) of sniffs at food (Simon, Widdig & Weiß, bioRxiv 2022.08.08.503203). Within a project on the role of olfaction in this catarrhine primate, we here investigated the interplay between visual information and olfaction in two experimental feeding contexts. In experiment 1, we provided Barbary macaques with food or non-food items in either a visible or non-visible condition to assess i) if visibility affected the propensity to sniff at the setup during any stage of exploring it, ii) if visibility affected at what stage of the exploration monkeys used olfaction, and iii) the interplay between visual condition, olfaction and the (attempted) retrieval of items from the setup. We expected monkeys to be more likely to sniff, and use sniffing as the first type of inspection if visual information was absent. We further expected that they use olfactory information gained in the non-visible condition to inform subsequent behavior (i.e. attempting to retrieve the content or not). In experiment 2, we manipulated the visual appearance of a familiar food item, i.e. popcorn, with food dye to assess how the salience in visual information affected sniffing behavior. We expected that popcorn with unfamiliar visual appearance (i.e. dyed popcorn) would be sniffed more frequently than naturally colored popcorn. We initially also intended to assess whether sniffing and visual appearance jointly affected whether or not items would be discarded or eaten, but restrictions in the experimental design as well as a very low prevalence of discarded popcorn prevented us from explicitly testing this.

## METHODS

### Study population

The study was conducted at Affenberg Salem, Germany in February and March 2021. Affenberg Salem is home to ∼ 200 Barbary macaques in three naturally formed social groups that range freely year-round within a 20ha forested enclosure. The park is open to visitors from spring to autumn, and is an active site of research year-round. Visitors are restricted to a designated path, while researchers have access to the entire park area. As a result, the monkeys are well-habituated to human presence (see de Turckheim and Merz 1984 for details on the park). The monkeys feed on plants and insects found naturally within the park and are supplemented daily with fruit, vegetables and grain. Water is available ad libitum in ponds and water troughs throughout the park. Individuals are identifiable from natural markings such as facial pigmentation patterns, coat color and stature as well as an alphanumeric code tattooed on the inner thigh. Individual life histories, group memberships and maternal relatedness are known from near-daily monitoring by staff.

### Experiment I: transparent *vs.* opaque pipes

#### Setup

We experimentally presented the monkeys with different food and non-food items in a visible and non-visible condition at 12 sites throughout the park. Hence, experiments took place in a near-natural environment without spatial or temporal restrictions of access. For logistic reasons, we split the experiment into three blocks, during each of which four sites were active simultaneously. In each block, the four active sites were distributed across the home ranges of all three groups. Sites were set up in locations routinely used by the monkeys but not directly at the feeding areas.

At each site we fixed a grey drainage pipe (11cm diameter) with a 90° angle to the bottom of a tree. One, open end of the pipe faced upwards, the other to the right (when viewed from the front) as depicted in Fig. 1. The end pointing to the right was closed with a transparent or an opaque food storing can (made of hard plastic, 11cm diameter) using cable ties. The opaque storing can was identical to the transparent one but coated with a layer of grey duct tape (without appreciable odor to the human observers) on the inside. While we cannot fully exclude that monkeys may have been able to smell the tape inside the can, this should have no effect on whether or not monkeys sniffed, or when monkeys sniffed first (see description of response variables below), because any potential odor of the tape would only have become available after already having sniffed. Due to the angle and dimensions of the pipe, items placed into the can were not visible (as assessed separately by 3 different human experimenters) from the top opening of the pipe, but only through the walls of the can if this was transparent (Fig. 1). All cans were punctured three times at the bottom to allow the odors of the content to be perceivable not only from the top opening of the pipe but also when sniffing the bottom of the can (see Fig. 1). Pipes were fixed to trees with black belt straps (38mm width, polypropylene), with the opening at a height of 30-40cm. The dimension and height of the setup allowed monkeys to inspect the setup and also to grab into it from the top to retrieve items from the can.

**Figure 1:**
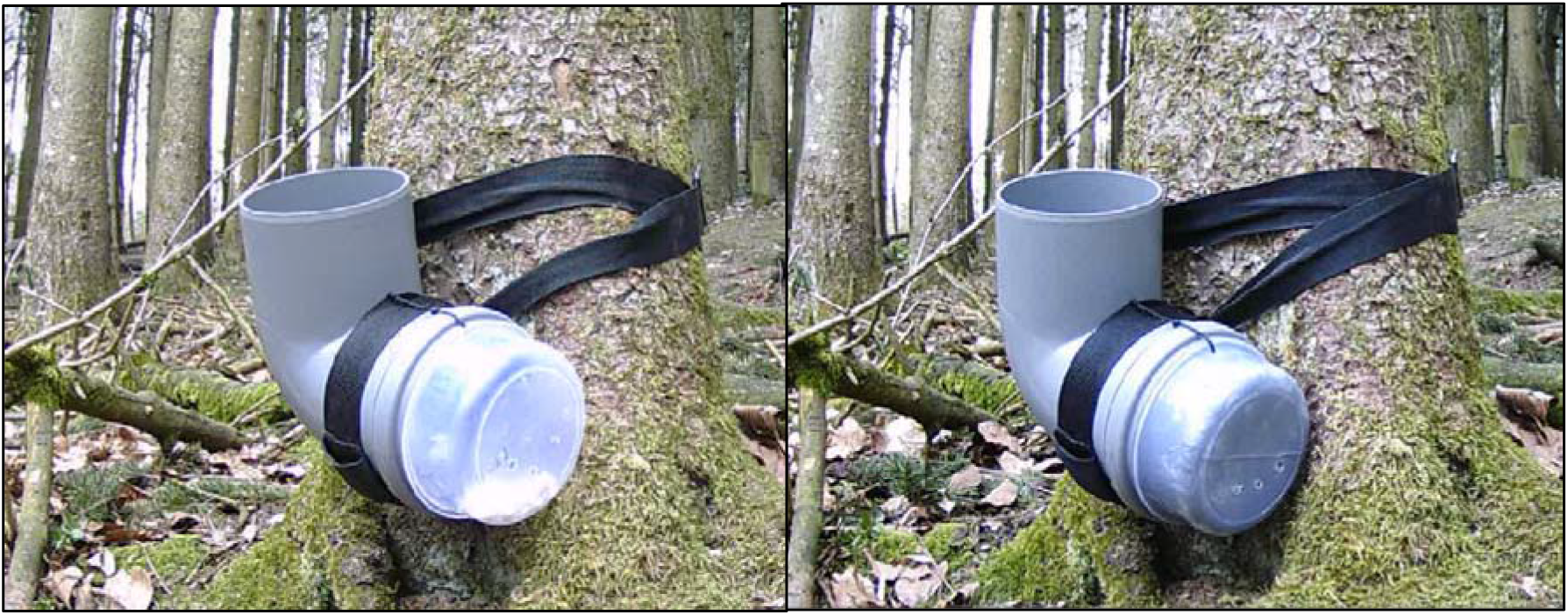
Setup for the pipe experiment with a transparent (left) or an opaque (right) storage can affixed to a drainage pipe. In the example shown, both cans are baited with popcorn, but the popcorn is only visible from the outside when the can is transparent. The setup is open at the top; and storage cans are punctured at the bottom to allow for olfactory investigation from either the top or the can.

At each site, approaches to and interactions with the setup were continuously monitored with two wildlife cameras (Crenova 4K 20 MP) mounted to neighboring trees to cover the setup from two different angles. Cameras were set to record a full HD video (avi format with 1920×1080p resolution) for 1min without sound whenever detecting movement with their motion detectors (see Supplement S1 for details). Cameras were equipped with a night mode which used infrared light, allowing for 24/7 monitoring.

#### Experimental design

At each site we first conducted a 2d habituation phase to familiarize the monkeys with the setup and its location. During the entire habituation phase a transparent can was attached to the pipe, and the pipe was filled with food items (3-6 pieces of peeled peanuts or popcorn) at random intervals 6-8 times per day.

The habituation phase was followed by a 7d test phase. In the test phase, the transparent can was exchanged with an opaque can every few hours and the can was filled with different food or non-food items or was pretended to be filled but left empty (see details below). At each site, the setup was filled 6x/d using the following semi-randomized manner: we divided daylight hours (7 a.m. to 7 p.m.) into six 2h time slots. In each slot, a given setup was filled once within the first hour of the slot, whereby we randomly determined the start time (0, 15, 30 or 45 min of the first hour) for filling the first setup and the order in which the four sites deployed simultaneously were visited. This ensured that the experiment covered the different daytimes evenly without making the visits to the respective setups easily predictable. In this way there further was at least one hour between consecutive visits to the same site, thereby increasing chances that one or several monkeys passed the setup while filled with a given content.

At each visit, the can was filled with one of 7 possible contents (5 types of contents, 2 of them in two different quantities as described below). To provide an incentive for exploring the setup with different senses rather than grabbing inside by default, we did not only fill the setup with food items but also non-food of similar size to the food items or nothing at all. As food items we either used pieces of peeled peanuts or popcorn, which were provided in two different quantities (3 or 6 pieces) to vary the intensity of their odor. As non-food items we either used 3 stones or 3 small pieces of Barbary macaque feces. We chose stones as presumably neutral, odorless items (i.e. neither particularly desirable nor undesirable for the monkeys). Stones with an approximate size of the peanut pieces were collected within the park area, washed with a neutral soap and water, and then rinsed with distilled water before use in the experiment. In contrast, feces were used as a potentially aversive item, as contact with feces may increase the risk of contracting infections and is avoided in numerous species (e.g. Case et al. 2020; Lopes et al. 2022). Feces were collected opportunistically whenever individually identified, adult monkeys were seen to defecate. Feces were stored in 30ml glass vials with screw top sealed with parafilm and were frozen at -20°C within 2h after collection. Feces were defrosted ∼30min before use and sized to approximately match the size of the peanut pieces. We used feces from both sexes but from other groups than the group using the area of the experimental site. As the experiment was conducted in a near-natural environment without spatial or temporal restrictions in access to the setup, monkeys could potentially observe the filling of the setup. To better disentangle whether a filling event or the items in the setup affected exploratory behavior, we therefore included a 7^th^ possible content, for which the can was pretended to be filled by reaching into it, but was left empty. All items were handled with disposable gloves, and the experimenter wore a medical mouth-nose cover. At each visit potential remains of the previous filling were removed, the inside of the setup was wiped first with 70% Ethanol (Carl Roth) and then with distilled water to remove residues of the ethanol before being refilled.

Over the course of the 7 test days, each of the 7 possible contents was presented once in the transparent and once in the opaque condition in three of the six 2h time slots: one of the first two, one of the middle two and one of the last two slots of the day (i.e. 7 contents x 2 conditions x 3 daytimes = 42 times). The order of these 42 filling events was randomized conditional on the following rules: a maximum of 3 consecutive fillings without edible items and no empty conditions twice in immediate succession at a given site (to keep the incentives up), and at least one but a maximum of three changes between transparent and opaque cans per day and site (to balance randomization with feasibility). This randomization procedure was done separately for each site, so that the order of conditions and contents was also randomized across sites.

#### Coding

We first screened all recorded videos and discarded the “false alarms”, in which the cameras were triggered by movement in the background or other animals than the monkeys (e.g. mice at night). We kept a total of 6570 videos in which one or more monkeys approached (i.e. interacted with or passed within 1m of) the setup. This number comprises the videos recording the same approach(es) from both angles by different cameras, whereby we only coded one of the simultaneously recorded videos if all behaviors were discernible from a single angle. It further comprises 181 videos recorded in the early morning hours after the end of the 7d test phase but before the setup was dismantled.

From the videos we scored each individual monkey’s approach. Videos could encompass multiple approaches per video or multiple, successive videos per approach, if the monkey remained at the setup for more than 1 minute. For each approach we scored from the video the date and start time as well as whether or not the individual looked, sniffed or grabbed into the setup (see sample videos in Supplements S2a-c). Looking was scored when the approaching animal oriented its eyes towards or into the opening or can of the setup. This was typically accompanied by a clearly noticeable movement and re-orientation of the head from a distance of less than 50cm, with the eyes closest to the setup while the snout and nose were pulled back towards the neck. Sniffing was scored when an animal placed its nose within 5cm of the opening or can, with the snout and nose stretched forward and placed closest to the setup. For both looking and sniffing we differentiated whether the behavior was directed at the opening of the pipe and/or can. Grabbing was defined as moving the hand (and arm) into the opening of the pipe so that at least the entire hand was inside the pipe. If a grab occurred, we also scored whether the monkey successfully retrieved anything from the setup. We scored any other interactions of the monkeys with the setup (e.g. climbing, tugging, biting into it) as ”other”, and further coded the order of the respective behaviors. The manipulation of the setup by the monkeys sometimes led to a slight shift in the pipe’s orientation. As a counterclockwise shift could potentially dislodge items from the can towards the bend, where it might have been visible from the opening irrespective of whether the can was transparent or opaque, we also scored the orientation of the pipe’s opening to statistically take into account the possibility that items may have become visible from the top opening (see Supplement S1 for details). All individual identities were scored from video by an observer with 2yrs experience in identifying the study animals. Behaviors were coded from video by a total of 6 different observers blind to the content of the setup, at least in the opaque condition (see Supplement S1 for details and inter-observer reliability).

Based on the experimental schedule and field notes, we determined information on condition (transparent or opaque), the exact time and content of the preceding filling for each approach at each site. By combining information on what was filled into the setup when, what was visible in the can on the video, what monkeys retrieved at a given visit to the setup and what was left in the setup at the next filling event, we were able to confidently deduce the actual content of the setup at the time of each individual approach.

For 7 of the 12 sites, the coded videos encompassed all monkey approaches to the setup (viewed from one or both angles) over the entire test phase. Hence, this also encompassed all approaches after (other) monkeys had already retrieved the items, which represents additional cases of an empty setup similar to when the setup was only pretended to be filled. At the remaining 5 sites, coding all approaches was not feasible because they were visited too frequently. For these sites we coded no more than the first 7 approaches after a given filling event and discarded the remaining videos (and thereby those approaches in which the setups were likely no longer baited). Overall, we analyzed 3590 videos, which corresponded to 1556 different approaches. All approaches took place during full daylight (98%) or late dawn (2%).

#### Statistical analysis

To assess the role of visibility on sniffing behavior as well as the consequences of olfactory inspection for further interactions of monkeys with the experimental setup, we constructed four generalized linear mixed models (GLMMs) in R (version 4.2.2, R Core Team 2022) using the function glmer in package lme4 (version 1.1-31, Bates et al. 2015). All 4 models were run with binomial error structure with logit link function. For these analyses, we only considered approaches in which the animals interacted with the setup (1431 of 1556 scored approaches, i.e. excluding those in which animals just passed by), all of which took place during daylight hours. To be able to account for repeated observations of the same individuals, we further used only those approaches for which we could identify the involved individual, i.e. 1399 approaches by 126 different individuals (which interacted with the setup 1-84 times). The sample sizes actually used differed between models for analytical reasons as described below.

As only food, but none of the other items were supplied in different quantities and a preliminary analysis suggested no differences in sniffing or grabbing behavior in relation to the quantity of items (Roderer 2021), we did not differentiate between quantities in the main analyses (models 1-4) to allow fitting all types of contents (peanuts, popcorn, stones, feces or nothing) in the same model. Details and results of a model investigating a potential effect of quantity on grabbing behavior when the setup contained peanuts or popcorn are presented in Supplement 1.

Model 1 was formulated to investigate the influence of the visibility condition (i.e. transparent or opaque can) on the propensity to sniff at the setup during any stage of exploring it. This model tested the prediction that visibility affects sniffing behavior. We fitted the model with sniff (yes/no) observed as response variable, using 1354 approaches which represented 853 cases with and 501 cases without sniffs by a total of 97 different individuals. As fixed effects test predictors, we modelled the visibility condition (transparent vs. opaque can), the actual content of the setup at the time of the approach (i.e. peanuts, popcorn, stones, feces or nothing) and the interaction between visibility condition and content to account for the possibility that different items were more or less likely to elicit sniffs in the transparent condition. We also fitted interactions between visibility condition and two control predictors: cohort (i.e. birth year of approaching individual) to account for age-dependent differences in the use of sensory modalities, and the number of days a given site was already active to account for a potential learning effect across time. As fixed effects control predictors, we further included sex (male/female), group (C, F or H), the experimental block (1-3, fitted as a covariate), the time (in min) since the last filling, the orientation of the pipe’s opening (see Supplement S1) and, to account for potential differences in activity patterns, the hour of the day. We further fitted the identity of the coder, the monkey ID, the date and the site as random intercepts to account for repeated measures and the random variation imposed by these grouping factors.

Model 2 used only data for the 7 sites for which all approaches had been coded (973 approaches by 80 different individuals of which 609 were with and 364 without sniffs). Coding of all approaches allowed us to also determine the time (min) since the previous approach to the setup by the approaching or other individuals, as well as to number the times the approaching individual had previously been at the setup. The motivation of this model was to take into account some of the individual information monkeys may have had available (from own experience or observing others at the setup) in addition to visibility of the content to guide their (olfactory) exploration of the setup. The model was constructed like model 1, but additionally included as fixed effects predictors the time (min) since approach by another monkey (delta other), time (min) since own previous approach (delta self) and n^th^ approach to the setup as well as the two-way interactions of these terms with the main test predictor, i.e. visibility condition. For the very first approach of an individual to a setup, we imputed the data by using the average time between approaches as time since own previous approach (delta self).

Model 3 was formulated to investigate whether the visibility condition affected if the monkeys used olfaction at the beginning of the exploration or later. Specifically, we expected the monkeys to sniff first when the content was not visible but later during the exploration if it was visible. Hence, this model used only those approaches in which the monkeys did sniff. As binary response variable, we fitted whether a sniff occurred first or later in the exploration sequence. Hence, 813 approaches by 71 individuals were used in this analysis. These represented 158 cases in which sniffs were the first behavior and 655 cases with sniffs at later stages of the exploration. Similar to model 1, we fitted the visibility condition, the content of the setup and their interaction, as well as the interaction between visibility condition and the control predictor day (i.e. number of days the site had already been active) as fixed effects test predictors. As fixed effects control predictors, we further included sex, cohort, group, block and the hour of the day. We further fitted the identity of the coder, the monkey ID, the date and the site as random intercepts.

Model 4 specifically investigated the interplay between visual condition, olfaction and the (attempted) retrieval of items from the setup. We expected that in the transparent condition vision will primarily guide the decision as to whether individuals would grab or not, while in the opaque condition olfaction should do so, and that in either case the decision would depend on the content. We therefore fitted grab (yes/no) as binary response variable, using the same 1354 cases as in model 1. These comprised 510 approaches with, and 844 without grabs. We fitted the visibility condition, the content of the setup, whether or not monkeys first sniffed (yes/no), and their 2-way and 3-way interactions as test predictors. In this model, ”sniff first” referred to the order of a sniff relative to the first grab to assess whether olfactory information may have guided the decision to grab. Hence, approaches with sniffs before the first grab, or sniffs not followed by a grab, would be denoted as ”yes”, and approaches with sniffs after the first grab, or without any sniffs as ”no”. As fixed effects control predictors, we further included sex, cohort, group, block, the day of the experiment, the time since the last filling of the setup and the hour of the day. We also fitted the interaction between orientation of the pipe and content to account for the possibility that food items (but not non-food items) having shifted into the bend could elicit more grabs. As in the other models, we further fitted the identity of the coder, the monkey ID, the date and the site as random intercepts.

#### General model procedures

To achieve reliable estimates for fixed effects predictors, we fitted all (model 1) or most (models 2-4) theoretically identifiable random slopes of fixed effects within the grouping factors (Schielzeth and Forstmeier 2009; Barr 2013, see Supplements S1 & S3). We did not fit any random correlations between random slopes terms to maintain an adequate balance between sample size and model complexity. We further excluded cases from the analyses that represented rare levels of grouping factors (i.e. fewer than 3 observations per level, see supplement S1) to facilitate accurate estimation of random variation.

For all models, we z-transformed covariates to a mean of 0 and an SD of 1 prior to fitting the model to facilitate convergence and interpretation of model estimates (Schielzeth 2010). To facilitate comparisons between studies, means and SDs of the untransformed covariates are provided in supplement S4. As the time since the last filling (delta fill) and since approach by another monkey (delta other) were strongly right-skewed, we log-transformed these variables after adding the constant 1 (to avoid 0-values) to improve model fitting. In these cases, the log-transformation was performed first and the z-transformation last. Furthermore, we manually dummy-coded categorical fixed effects predictors if involved in model interactions to be able to include their respective random slopes.

We checked for collinearity between predictors by using the function vif of the package car (version 3.1-1, Fox & Weisberg 2019). There were no issues with collinearity in any of the models (largest VIF in model 1: 1.15, model 2: 1.36, model 3: 1.25, model 4: 1.32). We further assessed the stability of models 1-3 by excluding each grouping factor level one at a time and comparing model estimates, using a function provided by R. Mundry. With a few exceptions (see Supplement S1), model estimates were generally stable. Because of the long computation time, we did not assess model stability for model 4 (estimated to take >60 days).

For inference, we compared the full models comprising all predictors with the corresponding null models, which excluded the test predictors but included all control predictors and all random intercepts and slopes. Hence, full-null model comparisons are indicative of whether one or more test predictors, but not whether control predictors significantly affect the response. Full-null model comparisons were conducted with Likelihood Ratio Tests (LRTs) using the R function anova. Only if the full-null model comparison was significant (p < 0.05), we assessed significance of the individual (test and control) predictors with LRTs using the function drop1. In case of non-significant full-null model comparisons we assessed significance of control predictors using Wald statistics provided by the glmer function, but did not consider results of test predictors. Accordingly, in contrast to test predictors, results of individual control predictors are not controlled for cryptic multiple testing. If a model contained non-significant interactions involving test predictors, we excluded the non-significant interactions from the model to facilitate interpretation of the main terms. For these models we present final models (after removal of non-significant interactions) in the main text and full models (including non-significant interactions) in supplement S5.

### Experiment II: colored popcorn

In the second experiment, conducted after the pipe experiment, we manipulated the visual appearance of a familiar food item, i.e. popcorn. This experiment was conducted with 32 adult focal animals of both sexes (16 males with ages of 7 to 21 years, 16 females with ages of 6 to 21 years) from two of the three study groups. All focal animals were already habituated to retrieving peanuts from a shallow plastic container in the course of another study.

With each of these individuals, we conducted two sessions (with 1-3d between sessions), in which popcorn was offered in a grey plastic container (the socket plug of a drainage pipe with 7.5cm diameter and 3 cm height) fixed to a horizontal branch or wooden rail with a belt strap. Each session consisted of a warm-up and a test trial. In the warm-up trial, a single piece of un-manipulated, white popcorn was offered in the container. If the animal participated in the warm-up trial, the container was filled with 4 pieces of popcorn simultaneously, 2 of which were white and the other 2 blue. Blue popcorn was obtained by dissolving a drop of viscous food dye (Dr. Oetker) in the same amount of water. The diluted dye was soaked up with a paper towel and softly dabbed over the entire surface of the popcorn, then left to dry. As we were unable to obtain white food coloring of the same brand, the white pieces of popcorn were left unmanipulated. In blind tastings of dyed and undyed popcorn, three experimenters (each tasting 3 pieces) were not able to distinguish between dyed and undyed popcorn, but we cannot exclude minor differences in smell or taste resulting from the dying process. To avoid interference by other monkeys, sessions were conducted when the respective focal animal had voluntarily separated from group members and was either out of sight or was at least 3m away from other monkeys and all individuals in the vicinity were lower-ranking than the focal animal. All sessions were conducted by the same experimenter and recorded with a handheld video camera (Panasonic HC-V180), with the experimenter commenting the observed behaviors and colors of the retrieved popcorn pieces while recording. All 32 focal animals could be successfully tested in this manner.

From the videos, we scored for each session the date and time, and for each piece of popcorn the order of retrieval, its color, whether or not it was sniffed at and/or eaten. All videos were coded by the same observer.

#### Statistical analysis

We analyzed the effect of popcorn color on sniffing behavior by fitting a GLMM with sniff (yes/no) as the binomial response variable with logit link. The unit of analysis was each single piece of popcorn offered to the 32 focal individuals in the test trials (N = 32 individuals x 4 pieces per session x 2 sessions = 256). As fixed effects test predictor, we fitted the color of the popcorn (white/blue). To account for the possibility that monkeys might get used to the novel coloration over the course of the experiment, we further fitted the interaction between color and the n^th^ piece of popcorn retrieved (continuous variable from 1 to 8). We controlled for sex (male/female), cohort, hour of the day and group (C or F) by fitting these terms as fixed effects control predictors, and for monkey ID and date by fitting them as random intercepts. We further fitted all identifiable random slopes of fixed effects predictors within ID and date (see Supplement S3), but no random correlations, as this caused non-convergence of the model. Checks of model assumptions and stability as well as determination of p-values for the full model and individual predictors were conducted as described in experiment 1. There were no issues with collinearity (largest VIF 1.31). Estimates for the effect of n^th^ piece of popcorn were rather wide and overlapped 0, but the model was stable with respect to the test predictor and the other control predictors.

##### Ethical note

Experiments were approved by Affenberg Salem. As all procedures were non-invasive and participation was voluntary, we required no further animal ethics approval.

## RESULTS

### Experiment 1: transparent vs. opaque pipes

We first addressed if visibility affected the propensity to sniff at the setup during any stage of exploring. In the 1399 approaches with interactions by identified individuals, monkeys sniffed in 439 (59.1%) of 743 approaches when the transparent can was attached to the pipe, and in 443 (67.5%) of 656 approaches when the can was opaque. However, although the proportion of sniffs in the opaque condition was slightly higher, the visibility condition had no significant impact on the propensity to sniff when the other predictors were controlled for (Model 1 on 1354 approaches by 97 individuals: full-null model comparison, LRT, χ2 = 11.747, df = 11, P = 0.383, see Tab. 1). This remained qualitatively the same when accounting for potential additional information on the contents from own or others’ prior approaches to the setup (Model 2 on 973 approaches by 80 individuals: full-null model comparison, LRT, χ2 = 17.807, df = 17, P = 0.401, see Tab. 1). While not addressed by the full-null model comparison, model results were suggestive of a significant effect of certain control predictors, namely cohort, with younger individuals being more likely to sniff than older ones in Model 1 (Wald test, z = 3.032, P = 0.002) and an interaction between cohort and visibility condition in Model 2 (Wald test, z = 2.015, P = 0.044), although the latter result was not very stable across models with different random level effects included (see details on model stability in the methods).

**Table 1:** Estimates of the GLMMs investigating the propensity to sniff for all experimental sites (Model 1, N = 1354 approaches) and for fully-coded experimental sites only (Model 2, N = 973 approaches).

|  | Model 1 |  | Model 2 |  |
| --- | --- | --- | --- | --- |
| *Term* | *Estimate* | *SE* | *Estimate* | *SE* |
| Intercept | 1.137 | 0.407 | 1.156 | 0.577 |
| Content |  |  |  |  |
| (feces) | 0.119 | 0.411 | 0.000 | 0.541 |
| (nothing) | -0.301 | 0.347 | -0.417 | 0.484 |
| (popcorn) | -0.664 | 0.406 | -1.085 | 0.523 |
| (stones) | -0.384 | 0.415 | -0.580 | 0.544 |
| condition (transparent) | -0.485 | 0.426 | -0.572 | 0.599 |
| delta_other |  |  | -0.117 | 0.125 |
| delta_self |  |  | -0.222 | 0.138 |
| n<sup>th</sup>_approach |  |  | -0.054 | 0.302 |
| Cohort | 0.328 | 0.108 | 0.287 | 0.129 |
| Day | 0.131 | 0.164 | 0.351 | 0.150 |
| sex (male) | -0.128 | 0.179 | -0.045 | 0.200 |
| Group |  |  |  |  |
| (F) | -0.114 | 0.265 | 0.185 | 0.410 |
| (H) | 0.164 | 0.327 | 0.125 | 0.578 |
| Block | -0.079 | 0.102 | -0.084 | 0.140 |
| delta_fill | -0.159 | 0.124 | -0.128 | 0.121 |
| Orientation | -0.081 | 0.071 | 0.008 | 0.081 |
| hour of day | -0.102 | 0.080 | -0.165 | 0.101 |
| content:condition |  |  |  |  |
| (feces:transparent) | -0.138 | 0.547 | -0.108 | 0.725 |
| (nothing:transparent) | 0.148 | 0.461 | 0.318 | 0.621 |
| (popcorn:transparent) | 0.691 | 0.584 | 0.927 | 0.836 |
| (stones:transparent) | 0.304 | 0.554 | 0.190 | 0.745 |
| condition (transparent):cohort | 0.172 | 0.136 | 0.351 | 0.174 |
| condition (transparent):day | -0.131 | 0.149 | -0.245 | 0.227 |
| condition (transparent):delta_other |  |  | 0.271 | 0.183 |
| condition (transparent):delta_self |  |  | 0.232 | 0.195 |
| condition (transparent):n<sup>th</sup>_approach |  |  | 0.120 | 0.190 |

Values in parenthesis indicate levels relative to the respective reference level (content peanuts, condition opaque, sex female, group C). SE = standard error. P-values not shown because full-null model comparisons were non-significant.

We further assessed, if visibility affected at what stage of the exploration monkeys used olfaction. n the 882 approaches during which the monkeys were observed to sniff, sniffing was the very first behavior directed towards the setup in 172 (19.5%) of cases, while the majority of sniffs occurred at later stages of exploring the setup. More specifically, monkeys inspected the setup first by sniffing in 76 (17.3%) of 439 cases in the transparent condition, and in 96 (21.7%) of 443 cases in the opaque condition, whereby both content and visibility condition significantly affected the propensity to sniff first (Model 3 on 815 approaches by 71 individuals, full-null model comparison LRT, χ2 = 24.782, df = 10, P = 0.006, see Tab. 2). Specifically, monkeys were slightly more likely to sniff first if the can was opaque, irrespective of content, and were more likely to sniff first when non-food items or nothing was in the pipe (Fig. 2).

**Figure 2:**
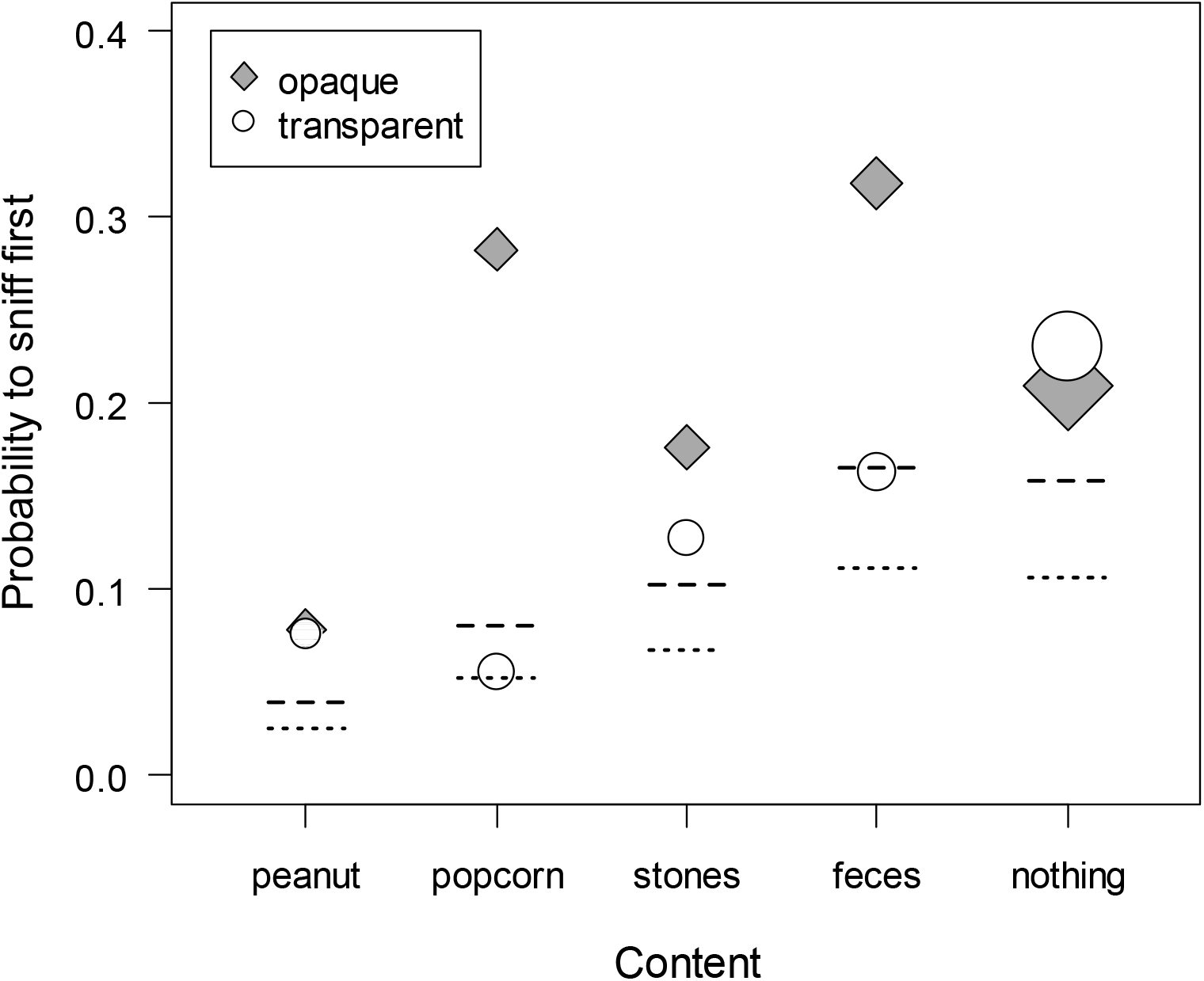
Proportion of approaches with sniffs in which the first interaction with the setup was a sniff depending on content and visibility condition (white symbols: transparent, grey symbols: opaque). Symbols are scaled by sample size (range 38 – 205 approaches with sniffs). Lines depict model estimates (dashed: opaque, dotted: transparent condition) when all other predictors are at their average.

**Table 2:** Estimates of the GLMM investigating the propensity to sniff first (N = 882 approaches.

| *term* | *Estimate* | *SE* | $\chi^2$ | *df* | *P* |
| --- | --- | --- | --- | --- | --- |
| Intercept | -3.061 | 0.787 |  |  |  |
| content |  |  | 14.061 | 4 | **0.007** |
| (feces) | 1.582 | 0.536 |  |  |  |
| (nothing) | 1.529 | 0.507 |  |  |  |
| (popcorn) | 0.766 | 0.574 |  |  |  |
| (stones) | 1.025 | 0.566 |  |  |  |
| condition (transparent) | -0.458 | 0.209 | 4.213 | 1 | **0.040** |
| cohort | -0.283 | 0.185 | 1.735 | 1 | 0.188 |
| day | 0.101 | 0.141 | 0.482 | 1 | 0.487 |
| sex (male) | -0.408 | 0.231 | 2.933 | 1 | 0.087 |
| group |  |  | 3.569 | 2 | 0.168 |
| (F) | -0.013 | 0.354 |  |  |  |
| (H) | 0.622 | 0.409 |  |  |  |
| block | -0.209 | 0.150 | 1.837 | 1 | 0.175 |
| hour of day | 0.084 | 0.163 | 0.281 | 1 | 0.596 |

Values in parenthesis indicate levels relative to the respective reference level (content peanuts, condition opaque, sex female, group C). SE = standard error. $\chi^2$ and P values are derived from LRTs to determine the significance of the individual test predictors. Significant effects are marked in bold.

Finally, we investigated the interplay between visual condition, olfaction and the (attempted) retrieval of items from the setup. In the 1399 approaches with interactions of identified individuals, the monkeys grabbed into the setup in 516 (36.9%) of cases. Whether or not the monkeys did grab was affected by the content, the visibility condition and whether or not the monkeys had sniffed at the setup (Model 4 on 1354 approaches by 97 individuals: full-null model comparison, LRT, χ2 = 94.424, df = 23, P < 0.001). In particular, the effect of the visibility condition depended on the content of the setup (Tab. 3). Specifically, monkeys showed a similar propensity to grab for food items irrespective of the visibility condition, but a reduced propensity to grab for non-food items or into an empty pipe in the transparent versus the opaque condition (Fig. 3). When the monkeys did grab, they actually retrieved peanuts and popcorn in almost all instances of grabbing (peanuts: 95 out of 96 times, popcorn 115 out of 119 times), but only in half of the 80 cases in which they grabbed stones and never in the 24 grabs when feces were inside the can. Sniffing generally decreased the propensity to grab into the setup (Tab. 3). Although this decrease appeared to be more pronounced in the opaque condition (Fig. 3), the interaction between sniffing and visibility condition or content was not significant. Furthermore, the propensity to grab inside was higher in younger animals and later in the experiment (Tab. 3). The quantity of the provided food items did not significantly affect grabbing behavior (Supplement S1).

**Figure 3:**
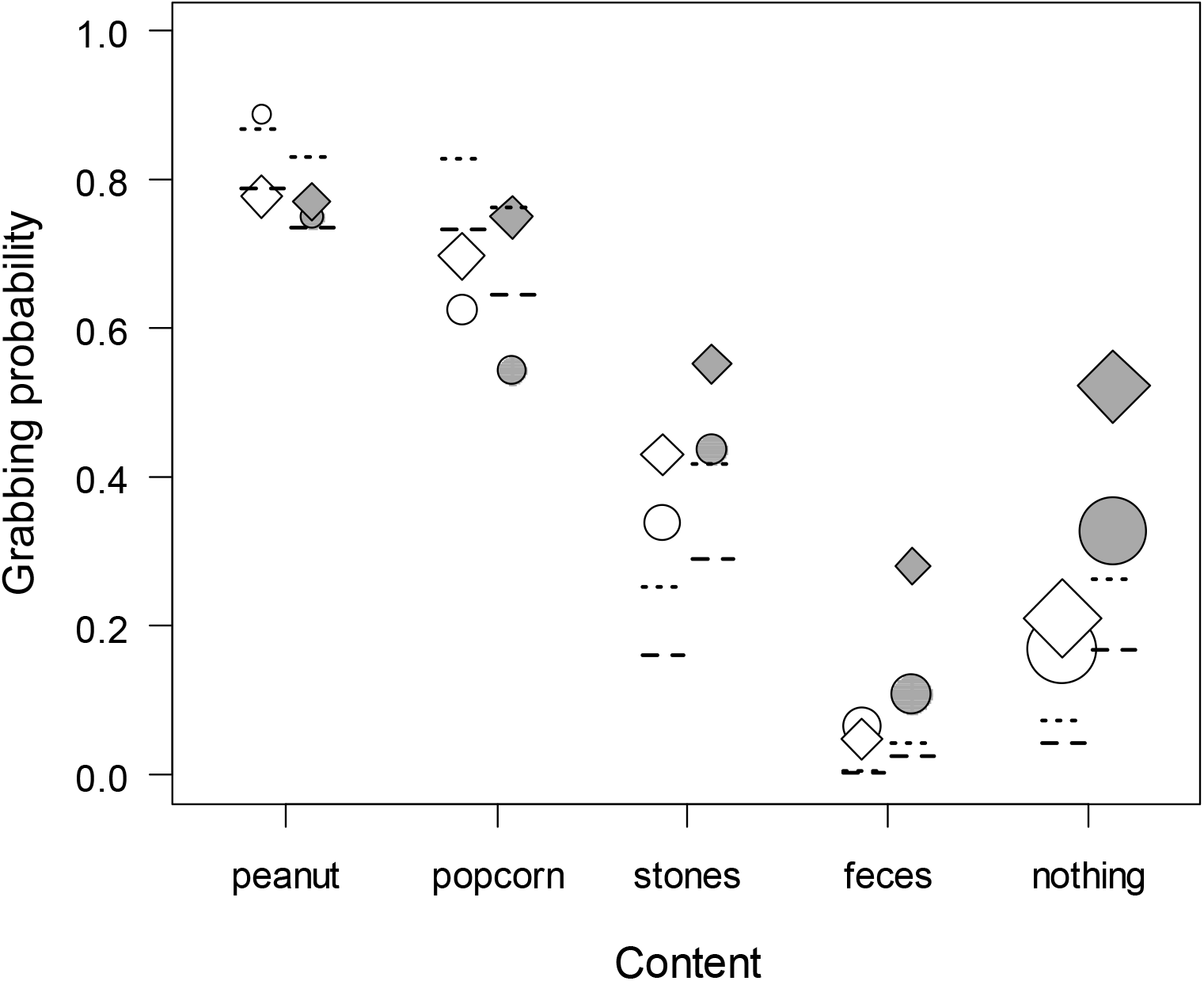
Proportion of grabs into the setup depending on content, visibility condition (white symbols: transparent, grey symbols: opaque) and whether or not monkeys had sniffed first (yes: circles, no: diamonds). Symbols are scaled by sample size (range 18 – 201). Lines depict model estimates (dashed: sniffed first, dotted: no prior sniff) when all other predictors are at their average.

**Table 3:**
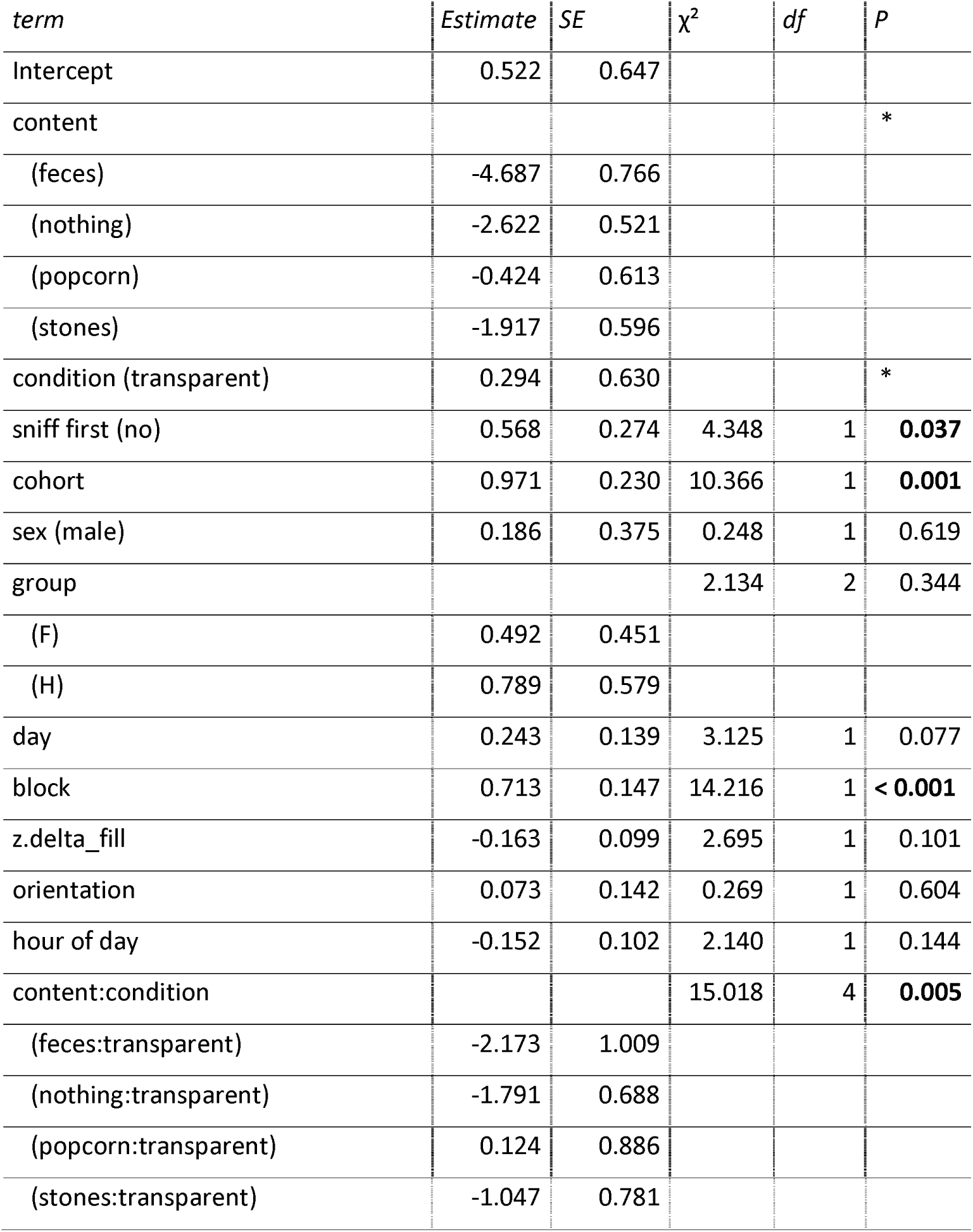
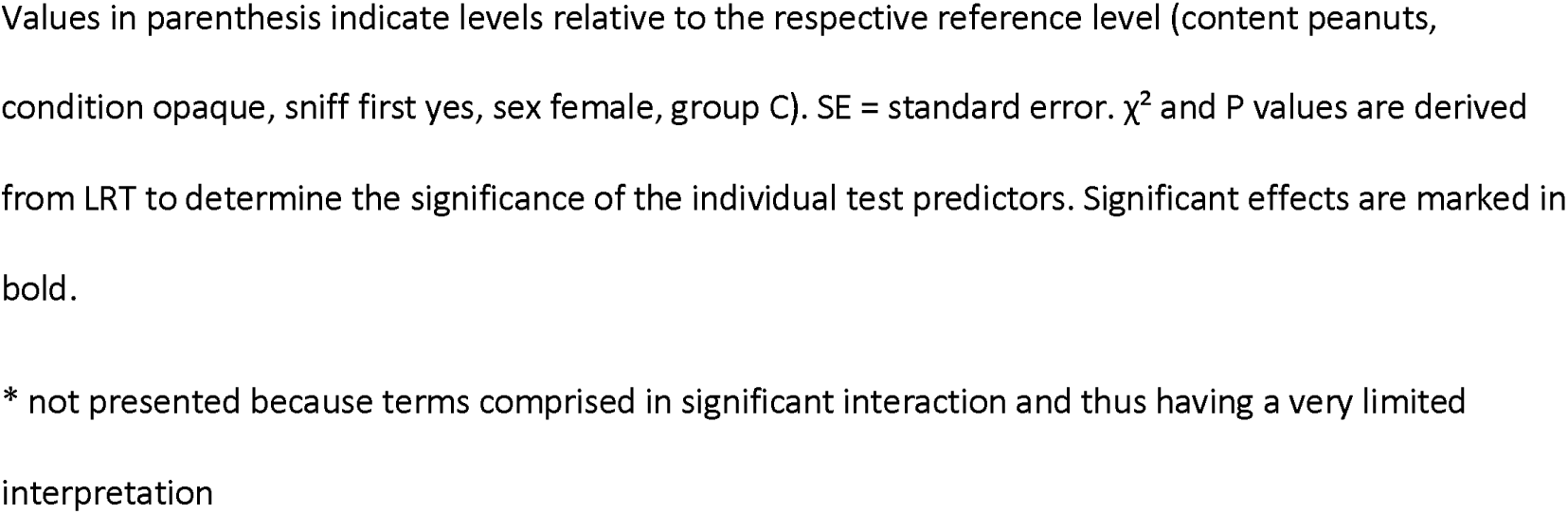
Estimates of the GLMM investigating the propensity to grab into the setup (N = 1354 approaches).

| *term* | *Estimate* | *SE* | $\chi^2$ | *df* | *P* |
| --- | --- | --- | --- | --- | --- |
| Intercept | 0.522 | 0.647 |  |  |  |
| content |  |  |  |  | * |
| (feces) | -4.687 | 0.766 |  |  |  |
| (nothing) | -2.622 | 0.521 |  |  |  |
| (popcorn) | -0.424 | 0.613 |  |  |  |
| (stones) | -1.917 | 0.596 |  |  |  |
| condition (transparent) | 0.294 | 0.630 |  |  | * |
| sniff first (no) | 0.568 | 0.274 | 4.348 | 1 | **0.037** |
| cohort | 0.971 | 0.230 | 10.366 | 1 | **0.001** |
| sex (male) | 0.186 | 0.375 | 0.248 | 1 | 0.619 |
| group |  |  | 2.134 | 2 | 0.344 |
| (F) | 0.492 | 0.451 |  |  |  |
| (H) | 0.789 | 0.579 |  |  |  |
| day | 0.243 | 0.139 | 3.125 | 1 | 0.077 |
| block | 0.713 | 0.147 | 14.216 | 1 | **< 0.001** |
| z.delta_fill | -0.163 | 0.099 | 2.695 | 1 | 0.101 |
| orientation | 0.073 | 0.142 | 0.269 | 1 | 0.604 |
| hour of day | -0.152 | 0.102 | 2.140 | 1 | 0.144 |
| content:condition |  |  | 15.018 | 4 | **0.005** |
| (feces:transparent) | -2.173 | 1.009 |  |  |  |
| (nothing:transparent) | -1.791 | 0.688 |  |  |  |
| (popcorn:transparent) | 0.124 | 0.886 |  |  |  |
| (stones:transparent) | -1.047 | 0.781 |  |  |  |

Values in parenthesis indicate levels relative to the respective reference level (content peanuts, condition opaque, sniff first yes, sex female, group C). SE = standard error. $\chi^2$ and P values are derived from LRT to determine the significance of the individual test predictors. Significant effects are marked in bold.
\* not presented because terms comprised in significant interaction and thus having a very limited interpretation

### Experiment 2: colored popcorn

The 32 focal animals retrieved all of the 256 pieces of popcorn and sniffed at 58 (22.7%) of them. The color of the popcorn had a clear impact on the probability of sniffing (full-null model comparison, LRT, χ2 = 10.77, df = 2, P = 0.005). More specifically, blue popcorn was sniffed at significantly more often (50 out of 128 times) than white one (8 out of 128 times), irrespective of how many pieces had already been retrieved (see Tab. 4). Furthermore, females generally sniffed at a higher proportion of popcorn than males (Tab.4, 5). Finally, the sniffing frequency appeared to decrease slightly over the course of the experiment irrespective of color, but this was only a statistical trend and not very robust (see details on model stability in the methods).

**Table 4:**
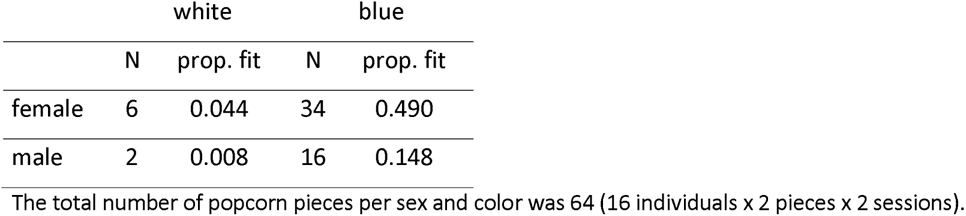
Number (N) and fitted values for proportion (prop. fit) of white and blue popcorn pieces sniffed by female and male Barbary macaques.

**Table 5:**
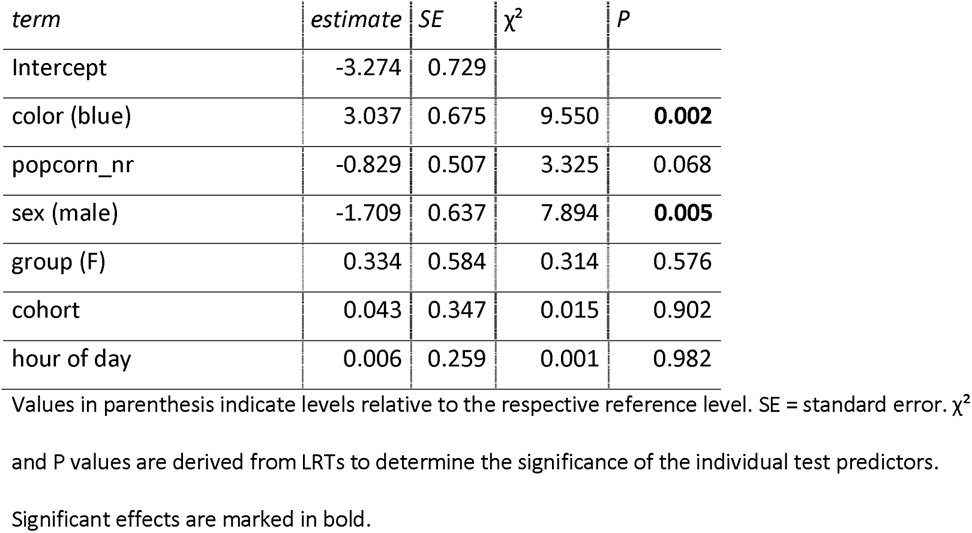
Estimates of the GLMM investigating the propensity to sniff at popcorn (N = 256 pieces of popcorn).

| *term* | *estimate* | *SE* | $\chi^2$ | *P* |
| --- | --- | --- | --- | --- |
| Intercept | -3.274 | 0.729 |  |  |
| color (blue) | 3.037 | 0.675 | 9.550 | **0.002** |
| popcorn_nr | -0.829 | 0.507 | 3.325 | 0.068 |
| sex (male) | -1.709 | 0.637 | 7.894 | **0.005** |
| group (F) | 0.334 | 0.584 | 0.314 | 0.576 |
| cohort | 0.043 | 0.347 | 0.015 | 0.902 |
| hour of day | 0.006 | 0.259 | 0.001 | 0.982 |

Values in parenthesis indicate levels relative to the respective reference level. SE = standard error. $\chi^2$ and P values are derived from LRTs to determine the significance of the individual test predictors. Significant effects are marked in bold.

The monkeys ate the vast majority of popcorn pieces (232) and only discarded 24 of them (23 blue, 1 white). They had sniffed at 13 of the discarded blue and at the one piece of white, discarded popcorn. Due to the low number of discarded pieces (especially white ones) and the lack of a positive control for the dye, we did not assess the interplay between visual and olfactory cues on the decision to eat or discard the popcorn statistically.

## DISCUSSION

The present study provides evidence that visual information affects the use of olfaction (i.e. sniffing) and subsequent exploratory behavior of Barbary macaques, whereby patterns differed depending on whether the presence or salience of visual information was manipulated in experimental feeding situations.

### Experiment 1: Effects of visual information on olfactory behavior (models 1-3)

Sniffing probabilities in experiment 1 were generally rather high and only little affected by the visibility condition: monkeys sniffed in more than half of the approaches even if the content was visible through the transparent can. Notably, sniffing occurred throughout all phases of interacting with the setup. While an initial assessment of the setup using olfaction was slightly more pronounced when cans were opaque than when they were transparent, the majority of sniffs in either visibility condition occurred after the monkeys had actively looked, grabbed or in other ways interacted with the setup. Hence, as suggested for various strepsirrhine and platyrrhine primates (reviewed in Nevo and Heymann 2015) and other mammals (e.g. swamp wallabies, *Wallabia bicolor*, Stutz et al. 2017), sniffing in Barbary macaques may be more relevant *after* an item is located through other sensory modalities (like vision), and thus may function primarily as quality assessment and food selection process rather than in the search for food.

The degree to which vision and olfaction interplay in food search and assessment has been associated with evolutionary adaptations to aspects such as diet or sensory abilities of foragers. In procellariform (”tube-nosed”) sea birds, for instance, differences in light exposure in early development have been related to the degree of dependence on visual and/or olfactory cues during foraging at sea, with burrow-nesting species depending more on olfactory and surface-nesting species more on visual or multimodal cues (Van Buskirk and Nevitt 2008). Similarly, the sensory cues provided by food items and the sensory capabilities of individuals may interact in an intricate manner in primate food selection. In spider monkeys, sniffing decreased from 40-45% in fruit species with rather cryptic coloration to 0 for the visually most conspicuous fruit species, irrespective of individual color vision abilities (Hiramatsu et al. 2009). In contrast, Melin et al. (2019) found that dichromatic capuchin monkeys sniffed more at fruits than trichromatic ones irrespective of the conspicuity of the visual cues provided by the fruits, although chromacy-related differences in sniffing rates varied greatly between fruit species and the overall difference was small (8% vs. 7% sniffing in di-vs. trichromatic individuals). Our study animals presumably all had highly similar visual capabilities with the one known exception of a subadult male with one blind eye from birth on. He sniffed in 70% of 37 approaches to the transparent and 73% of 22 approaches to the opaque setup and thus more frequently than the average. However, younger individuals were generally more likely to sniff (and grab into) the setup, and with just one case we are unable to assess whether the high sniffing rates were related to visual impairment, young age or just chance. Further studies incorporating a wider range of species are needed to address how aspects such as dynamic vs. stable differences in visual information, diet and other aspects of species ecology or individual differences in sensory capabilities affect to what extent vision affects olfactory behavior.

### Experiment 1: Effects of sensory information on grabbing for items (model 4)

Monkeys grabbed into the setup in a little more than one third of the approaches. As expected, whether or not monkeys grabbed was modulated by the content and its visibility. They were most likely to grab for peanuts and least for feces in either visibility condition, but the differences in grabbing between contents were less pronounced in the opaque condition. This suggests that the absence of visual information limited the monkeys’ ability to distinguish the contents of the setup, but also that they did have other sensory information than the items’ visibility available to guide the decision as to whether or not to grab inside. Our prediction that olfaction would mediate further exploration of the setup in the absence of visual information, however, was only partly supported. Results indeed indicated a significant effect of prior sniffing on the propensity to grab. Based on grabbing rates (see Fig. 3), this effect appeared to be more pronounced in the opaque condition, particularly when feces, nothing or popcorn was inside. However, statistically an interaction between sniffing, visibility condition and content could not be confirmed; model results rather suggest that monkeys that had sniffed the setup were generally less likely to grab inside irrespective of content or visibility condition. Although we scored well over 1000 approaches, the study design was complex and the number of actual approaches to a given content and condition as well as the (sniffing) behavior shown by the monkeys upon approach was not controllable by us, so that for particular combinations of predictors, cases may have still been too rare to detect statistically robust patterns. Another explanation could be that some residues of fecal odor remained in the setup even if other contents were inside, although we took great care to thoroughly clean the setup between fillings. Given the much lower propensity to grab for feces than for other items, this could explain a generally lower propensity to grab after sniffing irrespective of visibility condition and content, but if this was the case, we would have expected a much more pronounced decrease in grabs after sniffing than actually observed. We also consider it unlikely that a potential smell of the duct tape in the opaque condition affected grabbing, because sniffing the setup reduced grabbing irrespective of the visibility condition.

Notably, even if monkeys had not sniffed, grabbing rates differed between contents in the opaque condition (see Fig. 3), suggesting that the monkeys may have had other sources of information about the contents than anticipated. One possibility is that the odors of the contents may have been perceivable from a greater distance and without requiring the active sniffing movement that we scored, so that we may have underestimated the number of approaches in which olfactory information was available and used by monkeys. Although the majority of studies point towards primates primarily using olfaction for the close-up assessment of food items (Nevo and Heymann 2015), at least some species of strepsirrhines and platyrrhines appear to be able to detect and locate food sources from up to several meters distance using olfactory cues (Bicca-Marques and Garber 2004; Cunningham et al. 2021). Monkeys also sometimes pulled at the setup, which may have dislodged items from the tip of the can into a position visible from the top opening of the pipe. We statistically controlled for this possibility by including the orientation of the pipe into the models, but could not unambiguously determine for each approach whether or not monkeys could have seen the contents even in the opaque condition. Sound is unlikely to have provided salient information about the contents (other than maybe that there is something inside). Touch, on the other hand, has been suggested to be an important sense for evaluating food quality in primate feeding ecology (Dominy 2004; Veilleux et al. 2022). For instance, results by Dominy et al. (2016) suggest that the ingestion of green figs, *Ficus sansibarica*, by chimpanzees was elicited by fig toughness rather than their color or size. Along similar lines, the Barbary macaques in the present study may have grabbed inside the setup to gather tactile information about the item. The fact that the monkeys, in case of a grab, almost always retrieved food, but only half of stones and none of the feces supports the idea that monkeys may have also used tactile cues, but this possibility would need to be assessed systematically in a future study.

Finally, the food items used in the experiment are highly favored by the monkeys, and peanuts, in particular, are not routinely provided by the park. Hence, the incentive of gaining a favorite food item may have been higher than costs for an unsuccessful grab revealing no food or even an item like feces, so that at least some individuals may have pursued the strategy to grab inside as long as there was no obvious cue suggesting feces or other non-food content. Furthermore, monkeys not having encountered feces in the setup in previous approaches may have been more prone to grab inside irrespective of visual or odor cues to the content. However, taking into the account the prior setup experience of each individual would have required a much more detailed investigation including coding every single video, which was not feasible in the present study. Taken together, Barbary macaques appeared to depend more on visual than olfactory information to distinguish between items, although question marks remain as to whether or how visual and olfactory information interact to guide exploratory behavior in an experimental feeding context.

### Experiment 2

The second experiment addressed a different aspect of visual information. By presenting blue and white pieces of popcorn, visual information was constantly available but while size and shape were familiar, the color of dyed popcorn was not and thus represented a source of ambiguity. Indeed, monkeys sniffed popcorn in a novel (blue) color considerably more often than the familiar, white popcorn. This matches results in spider monkeys and squirrel monkeys, who used olfaction and other non-visual senses more when presented with novel food (i.e. entirely unfamiliar or with unfamiliar color or scent), while relying primarily on vision for assessing familiar food items (Laska et al. 2007). The unfamiliar visual appearance of the dyed popcorn also appeared to be particularly salient for sniffing behavior in Barbary macaques and elicited stronger changes in olfactory inspections than the presence or absence of visual cues in experiment 1.

Unfamiliar colors have also been shown to affect the acceptance and manipulation of food items (e.g. zebra finches: Kelly and Marples 2004; spider monkeys and squirrel monkeys: Laska et al. 2007). In our popcorn experiment, 23 of the 24 discarded pieces were dyed, but the monkeys consumed such a large proportion of the popcorn pieces (91%) overall that we did not have a sufficient number of discarded pieces to statistically assess how the interplay between visual information and sniffing behavior affected consumption. Furthermore, our inference about the role of visual appearance on food consumption was limited by the fact that we had no white food coloring of the same brand as the blue one available and thus compared dyed with undyed popcorn. Although different experimenters were not able to detect a difference in smell or taste of the dyed and undyed popcorn, we cannot exclude that the monkeys may have been able to perceive a difference. If that was the case though, this information should only have become available after sniffing or tasting the popcorn. Accordingly, a potential difference in the smell or taste of dyed popcorn should have been of little relevance for the question if an unusual color elicits olfactory investigation, but might have had an effect on the consumption of the popcorn. Follow-up studies should therefore incorporate different colors as well as combinations with odors (as done in e.g. Kelly and Marples 2004; Laska et al. 2007) to get a better understanding how unfamiliar visual appearance and olfactory information interact in catarrhine primate food choice.

## Conclusions

The present study provides evidence that Barbary macaques used visual and olfactory information to guide exploratory and feeding behavior in experimental feeding contexts. As a highly visually oriented species with acute color vision it comes as no surprise that the presence or absence as well as the salience of visual information clearly affected how monkeys responded to different food and non-food items. Monkeys also routinely used olfaction to inspect items, particularly if visual information was ambiguous (i.e. familiar food in an unfamiliar color), while the presence/absence of visual information had only minor effects on sniffing behavior. Olfactory inspection affected subsequent behavior to some degree, albeit less than vision. As such, this study adds to the increasingly comprehensive evidence that olfaction plays a prominent role in the lives of catarrhine primates, which were historically regarded to rely only little on the sense of smell. Importantly, by taking a multimodal approach we were able to show that the availability and salience of information in one sensory modality may affect the use of other sensory modalities, potentially leading to a complex, multisensory interplay affecting behavior. As also pointed out by various authors (e.g. Smith and Evans 2013; Leonard and Masek 2014; Valenta et al. 2015; Wilke et al. 2017; Veilleux et al. 2022), our results highlight the importance of a multimodal perspective, as examining single modalities may be insufficient to explain behavioral decisions. How such multisensory interactions guide behavior in a feeding or other contexts requires further studies and across a wider range of species to understand how (socio)ecological traits shape sensory ecology.

## ACKNOWLEDGEMENTS

We thank R. & M. Hilgartner and the staff of the Affenberg Salem for continuous support and E. Merz for permission to conduct this study. We further thank M. Roderer and M. Zschiesche for filling the setups and operating the wildlife cameras, and M. Roderer for performing the popcorn experiment. J. Kromp, N. Ritter, M. Roderer, L. Schmidt and H. Westphal helped with coding the videos. R. Mundry provided R functions for assessing model assumptions and stability.

