## Supplement S1 for "Influence of visual information on sniffing behavior in a routinely trichromatic primate"

**Supplemental Methods & Results**

Experiment 1: Setup

At each site, two wildlife cameras (Crenova 4K 20 MP) recorded 1min full HD videos whenever detecting movement. We set the break between consecutive videos of the same camera to the minimal possible setting of 5sec. Individuals interacting with the setup for longer than one minute were thus not consistently covered by a single camera. However, as the two cameras at a given site were affixed to different trees and monitored slightly different angles/ranges, monkeys came into their range at slightly different times and thus triggered the two cameras with an offset of a few seconds from each other, thereby allowing continuous monitoring of the setup by at least one of the cameras nonetheless. Pipes and cameras were oriented in a way to avoid strong backlight and a background with much activity (e.g. feeding area, visitor path) to improve image quality and reduce the number of false triggers of the cameras.

Experiment 1: Coding

*Pipe orientation:* To account for slight shifts in the pipe's orientation and the possibility that items may have become visible from the top opening, we scored the angle of the opening. We scored the orientation of the pipe from -5 to +5, with 0 resembling the opening pointing straight to the top. Positive numbers reflected the opening tilting to the right (i.e. no possibility of dislodging contents from the can), and negative numbers to the left (i.e. a possibility of items shifting towards the bend and thus being visible from the opening). The position of the opening was positive or neutral (0) in 83.9% of approaches used for analysis (see below), tilted marginally to the left (-1) in 15.2% of approaches and tilted more strongly to the left (-2 to -5) in only 0.9% of approaches.

*Interobserver reliability:* Videos were coded by a total of 6 different observers, of which four had previously worked with Barbary macaques at Affenberg Salem, while two had no prior experience with observing nonhuman primates. Before coding, pairs or trios of observers were trained together.

In a subset of 396 approaches, behavior was independently coded by two observers, which showed an inter-observer agreement of 87.4% for looking, 74.7% for sniffing and 94.2% for grabbing into the setup. To assess the reliability of identifying individuals, identities were coded by a second observer (with 6 months experience with the study population) in 70 approaches. The two observers agreed in their assessment of individual identities in 67 (95.7%) of these approaches.

<u>Experiment 1: General model procedures</u>

*Random slopes:* For model 1 we fitted all theoretically identifiable slopes. Overall, this model thus included a total of 75 estimated terms (fixed effects, random intercepts and random slopes).

Because of the lower sample size but higher number of predictors for model 2, we slightly simplified the random slope structure by fitting all theoretically identifiably random slopes for test predictors but only slopes for control predictors if predictors varied in at least two thirds of the levels of the respective grouping factor (see Supplement S2), resulting in a total of 92 estimated terms.

Similarly, for model 3 we fitted all theoretically identifiably random slopes for test predictors, but only slopes for control predictors if predictors varied in at least two thirds of the levels of the respective grouping factor (see Supplement S2). This model included a total of 61 estimated terms before and 40 after exclusion of non-significant interactions (see general procedures).

In model 4 we attempted to fit all theoretically identifiably random slopes, but this model did not converge. We therefore ran the model with a simplified slope structure containing only slopes of predictors varying in at least 3/4 of the levels of the respective grouping factor (but all the slopes for the main terms of test predictors, see Supplement S2). This model included a total of 105 estimated terms before and 71 after exclusion of non-significant interactions.

*Excluded cases:* To facilitate accurate estimation of random variation, we excluded 45 of the 1399 cases from models 1 and 4 that represented rare levels of grouping factors, which was the case for one date with only 2 scored approaches and 29 individual monkeys with one or two scored approaches. In the same manner 35 of the 1008 cases were excluded from model 2, and 69 of the 882 approaches with sniffs from model 3, all of which represented rare levels of date and ID.

*Model stability*: In model 1, estimates for the test predictors and some control predictors (in particular sex, group and block) were rather wide, but generally had the same sign or were close to 0, suggesting stable model results with respect to the sign of effects, but some uncertainty in the magnitude. This was also the case in model 2, with the exception of the interaction between visibility condition and cohort, which varied moderately between positive and negative values depending on the levels of random effects included in the model. In model 3, stability was estimated based on 111 models lacking one single level of a grouping factor. These estimates were generally stable with the exception of 4 out of 111 estimates for content and one out of 111 estimates for group, where the removal of a random effects level resulted in highly deviating estimates. In all 5 cases, deviations in the estimates were explicable by considerably smaller sample sizes due to the removal of the respective random effects level and/or complete separation, suggesting that the results overall were stable.

Experiment 1: Quantity of food items

*Model details:* To investigate whether the quantity of the provided food items affected the monkeys' propensity to grab for items, we used only approaches in which the setup was filled with either peanuts or popcorn. As in model 4, we fitted grab (yes/no) as binary response variable and the visibility condition and whether or not monkeys first sniffed (yes/no) as test predictors. As an additional test predictor we fitted the quantity (i.e. n items) in the setup at the time of each approach. We further fitted the two-way interactions between quantity and visibility condition, and between quantity and whether or not monkeys sniffed first to take into account the visual and olfactory information the monkeys may have had about the (quantity of) the contents. As control predictors we fitted the content of the setup (peanuts or popcorn) and control predictors that were significant in the main analysis (model 4), i.e. cohort and block. Because of the reduced sample size

(298 approaches, of which only 85 resulted in no grab) we did not fit any other predictors fitted in model 4, and did not exclude rare levels of grouping factors, which would have reduced sample size even further. Random terms, checks of model assumptions and model inference were handled in the same manner as for the other models.

*Results:*

In the 298 approaches to the setup while containing peanuts or popcorn in different quantities, the monkeys grabbed into the setup in 213 (71.5%) of cases. Whether or not the monkeys did grab was not affected by quantity of the food items, irrespective of whether monkeys could see or had sniffed the contents (298 approaches by 83 individuals: full-null model comparison, LRT, $\chi2 = 2.211$, df = 3, P = 0.53, Table S1).

Table S1: Estimates of the GLMM investigating the propensity to grab into the setup depending on quantity when peanuts or popcorn were inside (N = 298 approaches).

| **term** | **Estimate** | **SE** |
|---|---|---|
| Intercept | 1.483 | 0.588 |
| content (popcorn) | -0.683 | 0.422 |
| quantity | 0.529 | 0.457 |
| condition (transparent) | 0.271 | 0.457 |
| sniff first (no) | -0.127 | 0.533 |
| cohort | 0.942 | 0.319 |
| block | 1.063 | 0.297 |
| quantity:condition (transparent) | 0.002 | 0.449 |
| quantity:sniff first (no) | -0.302 | 0.504 |

Values in parenthesis indicate levels relative to the respective reference level (content peanuts, condition opaque, sniff first yes). SE = standard error. P-values not shown because full-null model comparison was non-significant.
